# The *Rhizoctonia solani* RsLysM and RsRlpA effector proteins contribute to virulence by suppressing chitin-triggered immunity and hypersensitive response

**DOI:** 10.1101/395582

**Authors:** Fredrik Dölfors, Louise Holmquist, Panagiotis N. Moschou, Christina Dixelius, Georgios Tzelepis

## Abstract

*Rhizoctonia* (in Greek “root-killer”) species and particularly *R. solani* attacks a broad range of plant species and crops. It belongs to Basidiomycota and is a soil-borne pathogen causing mainly damping-off diseases of seedlings and root rot, although it can infect plants in any stage. Despite the severity of this disease, many aspects in *R. solani* infection biology still remain to be elucidated. Here we investigated the role of two effector candidates, predicted from the genome of a *R. solani* AG2-2IIIB strain that uses sugar beet as a host. Gene expression analysis showed that genes encoding for a LysM effector and a rare lipoprotein-A-like protein (RsRlpA) were induced upon early infection stages. When heterologous expressed in *Cercospora beticola* the two genes contributed to virulence. The RsLysM effector showed chitin‐binding affinity and suppression of chitin-triggered immunity but could not protect hyphae from hydrolysis. The RsRlpA effector suppressed hypersensitive response in *Nicotiana benthamiana* leaves. Overall, this study provides us with valuable information on *R. solani* infection biology, implying that this organism relies on mechanisms similar to hemibiotrophic pathogens in order to establish a successful infection.

## Introduction

*Rhizoctonia solani* Kühn is an important soil-borne pathogen causing substantial yield losses in a wide range of crops including cereals, soybean, potato and sugar beet (Adams, 1988). It is responsible for damping-off disease in many hosts (Anderson *et al.*, 1982). It incites crown and root rot in sugar beet, leaf spot and root rot in tobacco, sheath blight in rice and black scurf in potato (Ceresini *et al.*, 2002; Bernardes-de-Assis *et al.*, 2009; Buhre *et al.*, 2009; Gonzalez *et al.*, 2011). The fungus produces persistent sclerotia, which are the major infestation source together with fungal-infested plant debris in the soil. *Rhizoctonia solani* commonly infects roots and hypocotyls but all types of plant organs can be colonized by the mycelia. The sexual stage (*Thanatephorus cuccumeris*) is extremely rare (Adams and Butler, 1983) and conidia formation is lacking. This pathogen is classified into different anastomosis groups (AGs) based on hyphal cell wall fusion between different isolates. Some AGs are further divided into subgroups based on host range, colony morphology, pathogenicity, zymogram patterns and other characteristics (Ogoshi *et al.*, 1987).

Despite the economic impact of *R. solani*, little is known about mechanisms behind fungal colonization and growth on host tissue. Hypothesis on gene function is difficult to test since *R. solani* is not amenable to molecular manipulations. A general view is that *R. solani* hyphae adhere to the host plant surface and enter the outer cell layers by the formation of penetration cushions (Gonzalez *et al.*, 2011), a process followed by the growth of invasive hyphae that ramify through the host tissue.

Secreted proteins promoting disease were first observed in human pathogenic bacteria (Preston, 2007). As more genomes of plant pathogens have been generated the prediction of secreted proteins or otherwise transported molecules have increased substantially. Evolution of effectors is a strategy by pathogens to manipulate plant defenses to establish a successful infection (Vleeshouwers and Oliver, 2014; Lo Presti *et al.*, 2015). A plethora of strategies are employed; effectors suppress and block early plant defense responses such as hypersensitive response (HR) (Hemetsberger *et al.*, 2012), stealth hyphae from recognition or protect them from plant chitinases (de Jonge and Thomma, 2009, van den Burg *et al.*, 2006) or promote necrosis (Qutob *et al.*, 2006). Key molecules acting in various pathways can also be hijacked impacting gene activation and hormones (Weiberg *et al.*, 2013; Ma and Ma, 2016). Many aspects on effector function during plant-microbe interactions remain unclear and host targets are most likely more than one. Genomic data are presently available from five different *R. solani* strains with cereal and dicot plant hosts (Cubeta *et al.*, 2014; Hane *et al.*, 2014; Wibberg *et al.*, 2013, 2016a, b; Zheng *et al.*, 2013). In an AG8 strain, attacking both monocots and dicots, a xylanase and a protease inhibitor I9 induced cell death when expressed in *Nicotiana benthamiana* (Anderson *et al.*, 2017). Similarly, cell death induction was also observed on rice, maize and soybean leaves, when purified candidate effectors from an AG1-1A strain, causing rice sheath blight, were evaluated (Zheng *et al.*, 2013).

Sugar beet is mainly attacked by the AG2-2IIIB strain. The disease can appear in different forms and with different symptoms; root and crown rot, damping off or as foliar blight (Bolton *et al.*, 2010). The demand for high yielding sugar beet cultivars with resistance to *R. solani* is increasing. Effects of a milder and more humid climate are seen in temperate regions influencing the length of crop seasons and simultaneously allow pathogens to multiply for longer periods. Moreover, the extent of *R. solani* AG2-2IIIB soil inoculum is expected to increase in regions particularly where sugar beet and maize are overlapping in the crop rotation schemes since maize can act as a host and thus propagate the pathogen (Buddemeyer *et al.*, 2004; Schulze *et al.*, 2016). The *R. solani* AG2-2IIIB strain has a predicted genome size of 56.02 Mb and 11,897 protein-encoding genes (Wibberg *et al.*, 2016a, b). Here we refined the dataset and identified 11 AG2-2IIIB-specific candidates. A rare lipoprotein-A‐ like (RsRlpA) protein and the chitin-binding lysin motif (RsLysM) effector were highly induced in infected sugar beet seedlings. In planta assays using *Cercospora beticola* as a heterologous expression system harboring the two *R. solani* genes showed increased virulence. The RsLysM effector was able to suppress chitin-triggered immunity, while the RsRlpA effector was able to suppress the hypersensitivity response (HR) upon transient expression on *N. benthamiana* leaves. These data give us valuable information regarding the different mechanisms that *R. solani* deploys upon the infection process and it seems that this pathogen does not only rely on necrosis-induced effectors.

## Materials and Methods

### Fungal isolates and sequence analysis

*Rhizoctonia solani* AG2-2IIIB isolate BBA 69670 (DSM 101808) (Wibberg *et al.*, 2016a) was used in this study. *R. solani* inoculum for soil infestation was prepared on media containing perlite, corn flour, and water. *Cercospora beticola* isolate Ty1 was grown on potato dextrose agar plates at 22°C in darkness and sporulation was induced on tomato growth extract medium. Small cysteine-rich and secreted proteins were retrieved from *R. solani* AG2-2IIIB genome (Wibberg *et al.*, 2016b). Presence of conserved domains was searched for by using the SMART 6.0 tool (Letunic *et al.*, 2009), followed by SignalP 4.1 for signal peptide prediction (Petersen *et al.*, 2011). The Phyre2 server was used for protein structure prediction (Kelley *et al.*, 2015).

### Quantitative reverse transcriptase - PCR (qRT-PCR)

Three-week old sugar beet plants were transplanted in soil infested with *R. solani* mycelia (10:1 ratio of fresh soil: inoculum). Total RNA was extracted from the plants at 4,5,6 and 7 days post inoculation (dpi) using the RNeasy Plant Mini Kit (Qiagen) according to manufacturer’s instructions, while *R. solani* mycelia grown in potato dextrose broth were used as fungal control. qRT-PCR was run and primers are listed in Table S1. Expression was normalized by *R. solani G3PDH* expression (Chamoun *et al.*, 2015) and relative expression values were calculated according to the 2^−ΔΔCt^ method (Livak and Schmittgen, 2001).

### *Cercospora beticola* overexpression strains and virulence assays

Overexpression vectors were designed using the In-Fusion HD cloning kit (Takara Bio). The RsLysM and RsRlpA sequences were PCR amplified from *R. solani* cDNA using high fidelity Phusion *Taq* polymerase (Thermo Scientific). The destination vector was the pRFHUE-eGFP, conferring resistance to hygromycin (Crespo-Sempere *et al.*, 2011). The *PgdpA*: effector construct was tagged with promoter with GFP in the C-terminus and transformed to *C. beticola* using an *Agrobacterium*-mediated protocol (Utermark and Karlovsky, 2008). Primers are listed in Table S1. Three independent colonies expressing the correct effector genes were selected for further analysis.

For virulence assays, leaves of 3-week-old sugar beet plants were inoculated with 10^5^ *C. beticola* conidia /ml H_2_0 from overexpressed, empty vector and wild-type (Ty1) strains. Lesion area was monitored at 7 dpi, and quantified using the ImageJ software version 1.51n (National Institute of Health, USA). Total genomic DNA was extracted from the diseased plant material using a CTAB-mediated protocol (Möller *et al.*, 1992). Fungal DNA was quantified using the *C. beticola* actin (*act*) reference gene and normalized with *B. vulgaris* elongation factor (*elf-1*) gene. Primers are listed in Table S1.

### Confocal microscopy and hypersensitive response assay

The RsRlpA effector sequence was cloned into pENTR/D-TOPO Gateway vector (Thermo Fisher Scientific), and entered to pGWB605 or pGWB660 binary vectors, tagged with C-terminus GFP or RFP fluorescence proteins respectively, driven by the 35S promoter and transformed into *Agrobacterium tumefaciencs* C58C1. Overnight cultures were used for Agro-infiltration on 4-week old *N. benthamiana* leaves. Protein cellular localization was monitored by using a Zeiss LSM 800 confocal microscope. For the hypersensitive response assay, the *RsRlpA* gene was entered to the pGWB602 binary vector without tag, driven by the 35S promoter, followed by Agro-infiltration in transgenic *N. benthamiana*, expressing the Cf-4 receptor protein from tomato plants (Joosten *et al.*, 1997). A HR resonce was triggered 24 hrs after Agro infiltration with the *Cladosporium fulvum* Avr4 effector protein in O.D 0.03 (Joosten *et al.*, 1994). The HR was evaluated using a scale from 0-3, ranging from no symptoms (0) to severe symptoms (3).

### Expression of RsLysM in *Pichia pastoris*

The RsLysM protein was cloned in the pPic9 expression vector with N-terminal His tag and transformed into the *P. pastoris* strain GS113. Primers are listed in Table S1A positive clone was cultured in a Bioflo 300 fermenter (Rooney *et al.*, 2005). His-tagged protein was purified using a Ni-NTA column (Qiagen) and final protein concentration was determined spectrophotometrically at 280 nm, and concentration analyzed by a Pierce BCA Protein assay kit (Thermo Fisher Scientific).

### Affinity precipitation, reactive oxygen species and hyphal protection assay

A chitin-binding assay of the RsLysM effector protein was performed. Pure protein was incubated with insoluble polysaccharides (crab shell chitin, chitosan, xylan or cellulose) and analyzed on SDS polyacrylamide gel (van de Burg *et al.*, 2006). For suppression of chitin-induced oxidative burst of reactive oxygen species (ROS) assay, the protocol by de Jonge *et al.* (2010) was used. Here leaf discs were inoculated with assay buffer with/without chitin oligomers (GlcNAc)_6_ and the suppression of ROS was tested using 10-20 μM RsLysM protein. For hyphal protection assay a protocol described previously by Kohler *et al.*, (2016) was used. Briefly, the protein was incubated with germinated conidia from *Trichoderma virens*. Then bacterial chitinase (Sigma) and 10U zymolyase (Zymo Research) were added. The Avr4 effector protein (5 μM) and BSA were used as positive and negative controls respectively. Images were taken 6 hrs post inoculation (hpi).

## Results

### *RsLysM* and *RsRlpA* are two effector candidates induced upon early-stage infection

In the previous genome analysis of the *R. solani* AG2-2IIIB isolate, 126 predicted secreted and cysteine-rich proteins were identified (Wibberg *et al.*, 2016b). To further reduce the list of effector-candidates only small proteins (<400 amino acids) were kept resulting in 61 predicted sequences. Eleven candidates appeared to be unique in this isolate. Analysis of sugar beet plants grown in *R. solani* (*Rs*) infested soil revealed elevated transcript levels of *RsLysM* (*RSOLAG22IIIB_4067*) and *RsRlpA* (*RSLAG22IIIB_8473*) genes already at 4 dpi suggesting an important role during *R. solani* plant host colonization (**Figure 1**). *RsRlpA* was predicted to encode a rare lipoprotein A (RlpA)-like domain protein, previously described in bacteria (Gerding *et al.*, 2009). Fungal LysM effectors are versatile proteins that occur in a plethora of species with divergent lifestyles. The general function of LysM molecules are to evade plant recognition but may also have other roles, for instance it is required for the appresorium function in *Colletotrichum higginsianum* (Takahara *et al.*, 2016).

**Figure 1.**
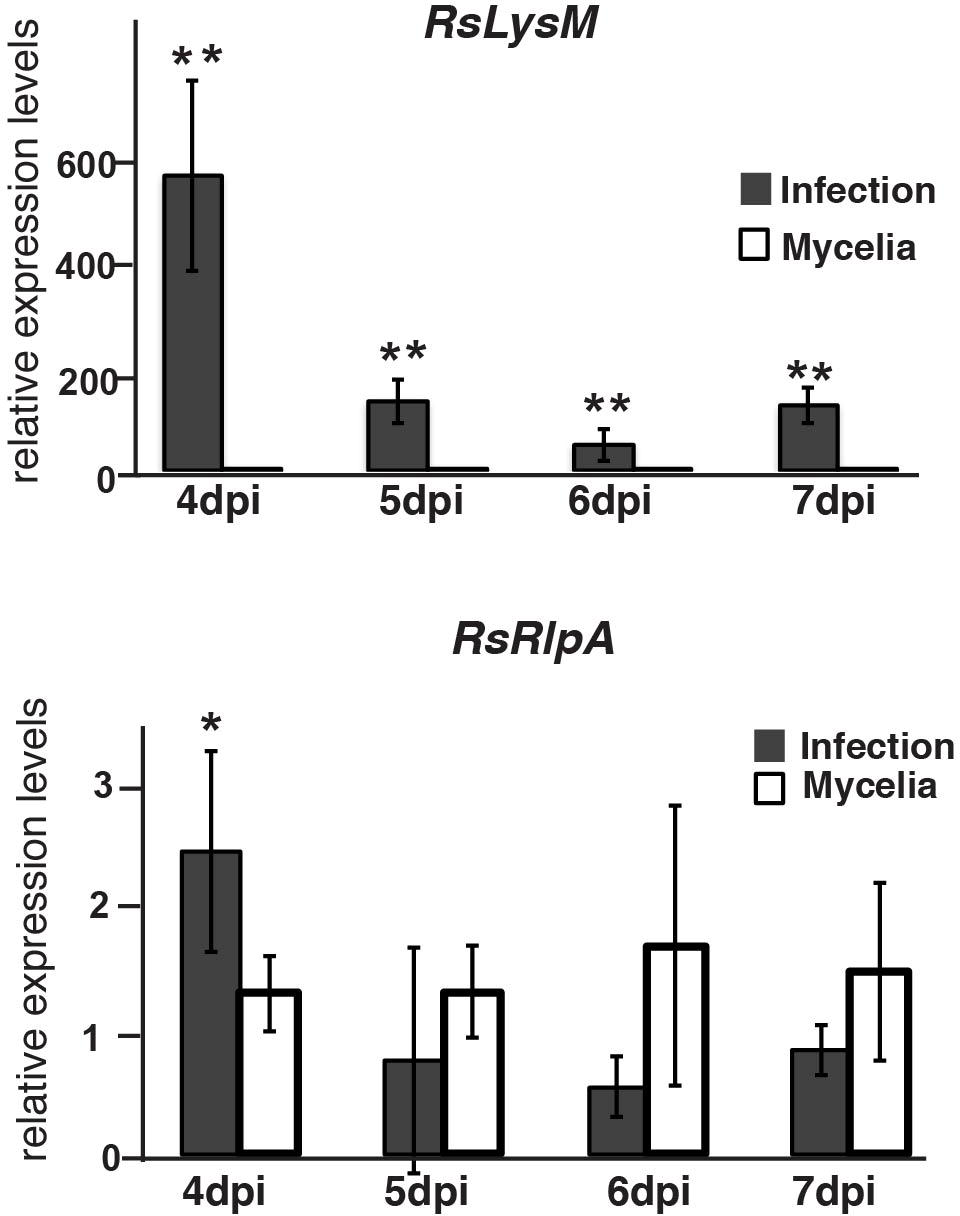
Relative transcript levels of the *Rhizoctonia solani* AG2-2I I I B genes *RsLysM* and *RsR/pA*. Sugar beet plants were grown in infested soil for 4,5,6, and 7 days before harvest for real-time qRT-PCR. *Rhizoctonia solani* mycelia was grown in PDB medium and used as comparison. The *G3PDH* gene was used as internal standard. Error bars represent SD based on at least three biological replicates. Asterisks(^*^ p value < 0.05, ^**^ p value < 0.01) indicate statistically significant differences between columns at the same time point according to Student’s T test.

### *RsLysM* and *RsRlpA* promote virulence upon heterologous expression in *Cercospora beticola*

The amino acid sequences of *RsLysM* and *RsRlpA* were analyzed for conserved domains. The SMART tool predicted two CBM50 modules (IPR002482; LysM peptidoglycan binding) in the RsLysM effector and a conserved double-psi beta-barrel (IPR036908; DPBB fold) in RsRlpA (**Figure 2A**). Structural prediction of RsRlpA using the Phyle2 resulted in: 39% homology to a bacterial cellulose binding protein, 34% to a papain-like inhibitor, 27% to expansin, 26% to kiwellin protein and 26% to barwin-like endoglucanase. Both RsLysM and RsRlpA proteins contain an ER signal peptide at the N-terminus, according to SingalP.

**Figure 2.**
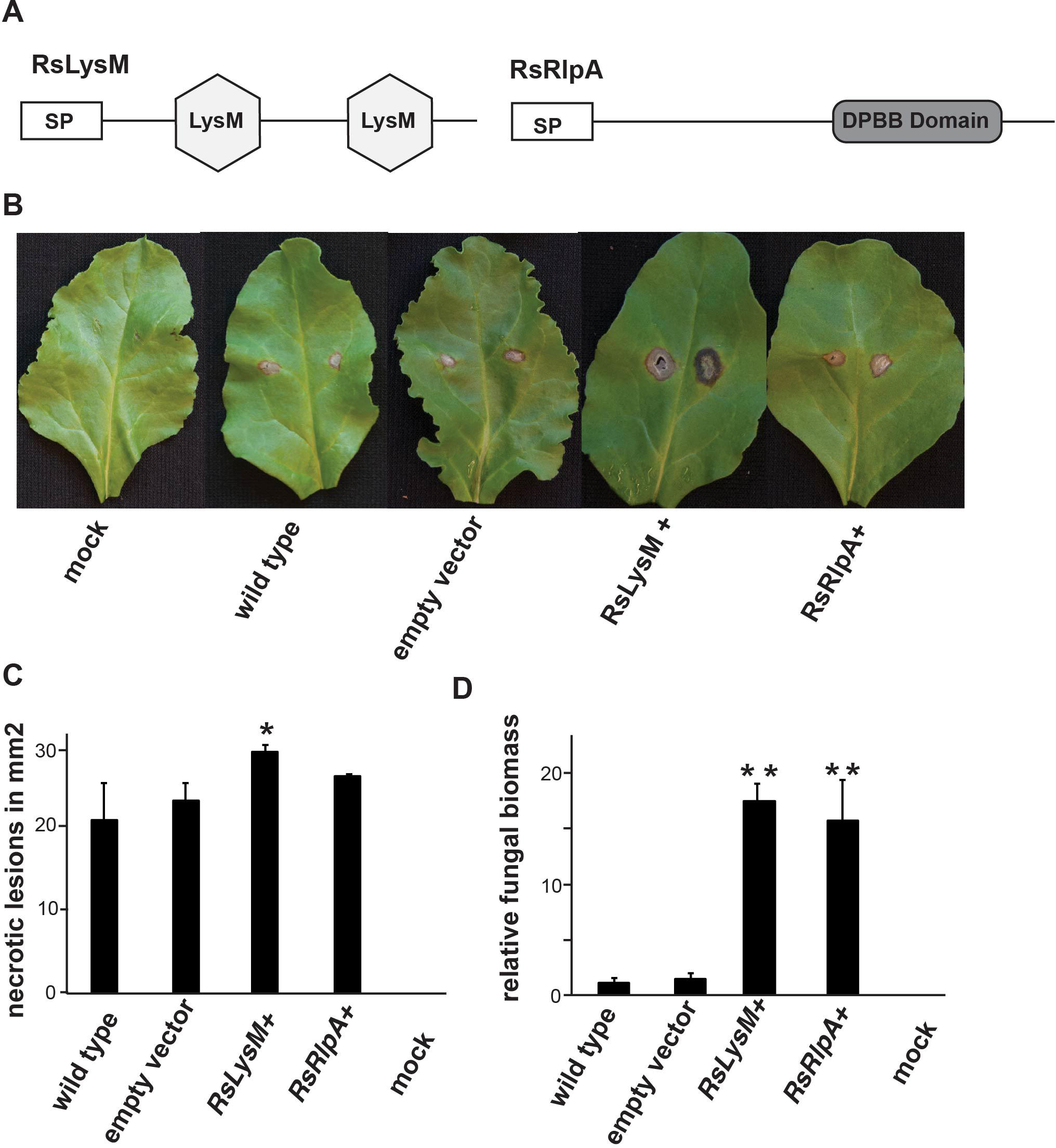
Assays of RslySM and RsRlpA candidate effectors. **(A)** Domain structure of RslySM and RsRlpA proteins. **(B)** Phenotypes on sugar beet leaves in response to C. beticola strains harboring RslySM and RsRlpA genes driven by the gdpA promoter, 7dpi. **(C)** Area of necrotic lesions on sugar beet leaves. **(D)** Fungal biomass quantification upon infection of sugar beet leaves. For quantitative PCR (qPCR), the C. beticola actin (act) gene was used. Data were normalized with the elongation factor gene (*e/f-1)* from *Beta vulgaris*. Data show the average of three independent overexpression strains each includes three biological replicates. Asterisks (^*^ p < 0.05, ^**^ p < 0.01) indicate statistic significant differences between the wild type, empty vector and overexpression *C. beticola* strains according to Fisher’s test. Error bars represent SD based on three independent strains.

To test if the two effector-candidates are involved in virulence, we heterologously expressed them in the sugar beet hemibiotroph fungal pathogen *C. beticola*. Using qRT-PCR we confirmed the expression of these genes in *RsRlpA*^+^ and *RsLysM*+ strains (**Figure S1**). Phenotypic analysis on these strains displayed no difference in morphology, growth rate or conidiation in comparison to wild-type (Ty1). The *RsLysM*+ *C. beticola* strains, displayed significantly larger necrotic lesions on sugar beet leaves, compared to wild type and strains where only the empty vector was inserted **(Figure 2B, C)**. In contrast, no clear phenotypic difference was observed in *RsRlpA*+ strains. Similarly, transient expression of RsRlpA in *N. benthamiana* did not induce cell death. The fungal biomass was significantly increased in comparison with wild-type and empty vector for both *RsLysM*+ and *RsRlpA*+ strains, further supporting the role of these candidate effectors in host colonization (**Figure 2D**). The localization of RsLysM and RsRlpA was checked in *C. beticola* fungal hyphae. Both candidate effectors were localized to the cell periphery, and RsLysM was particularly observed in hyphal tips (**Figure 3**).

**Figure 3.**
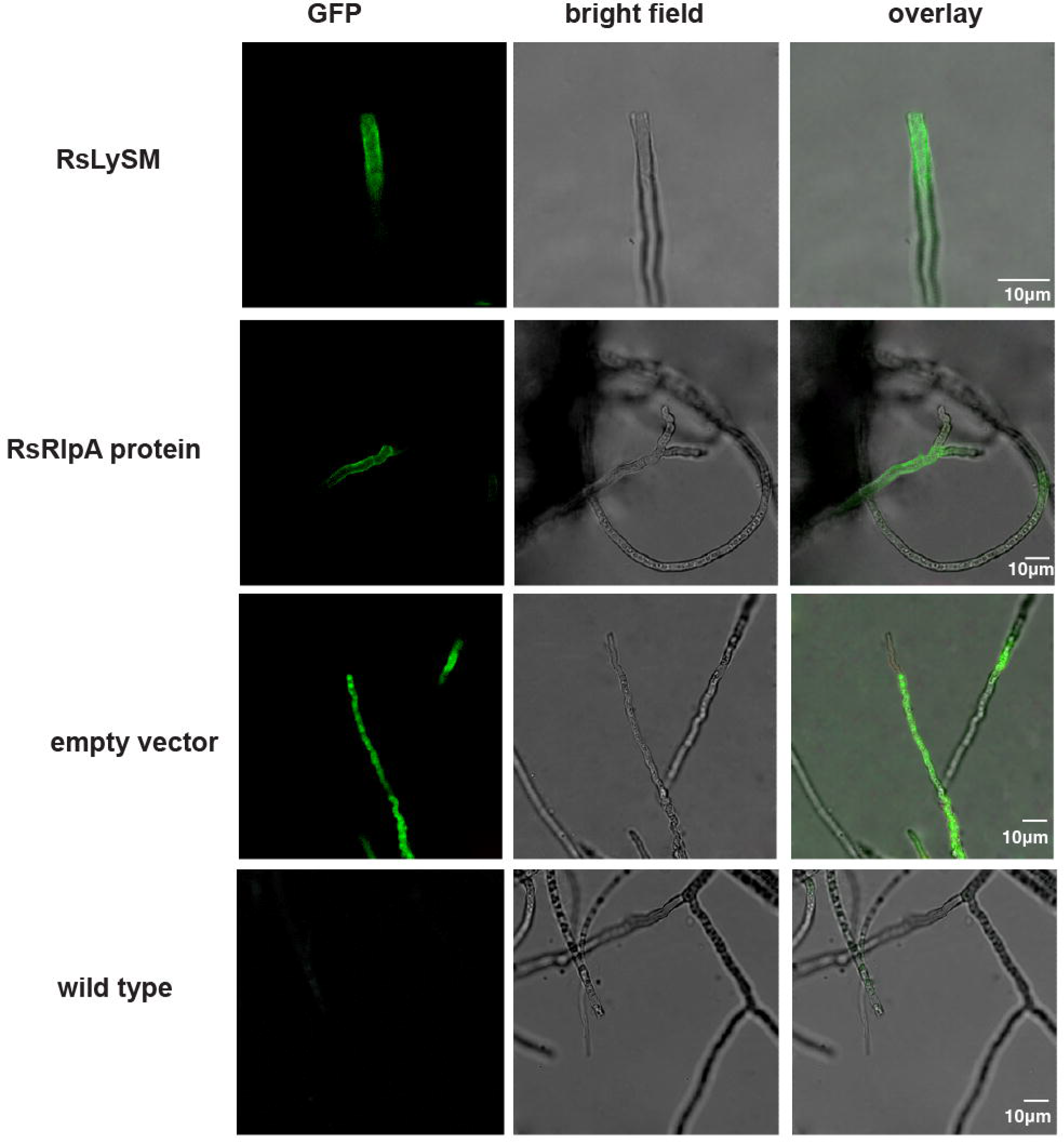
Live-cell imaging of GFP-tagged R. *solani* RslySM and RsRlpA effector candidates in C. *betico/a* hyphae. Live-cell imaging was performed with a laser-scanning confocal microscope with a sequential scanning mode, 7 days after inoculation on potato dextrose agar. GFP was excited at 488 nm and collected at 505-525 nm.

### The RsRlpA effector suppresses hypersensitive response and is mainly localized to the plant plasma membrane

Many plant pathogens rely on an initial short biotrophic stage to establish a successful infection and therefore secrete effectors to work as suppressors of HR (Stergiopoulos and de Wit, 2009). A hemibiotrophic stage has also been proposed for *R. solani* (Gonzalez *et al.*, 2011). Thus, the ability of RsRlpA effector to suppress HR was investigated by exploiting the Avr4-Cf4 interaction that leads to severe HR symptoms in *N. benthamiana*. Our data suggest that in leaf areas where RsRlpA had been previously Agro-infiltrated, reduced HR was observed as compared to the area where only *Avr4* was expressed (**Figure 4**). This observation suggests that RsRlpA potentially suppresses HR.

**Figure 4.**
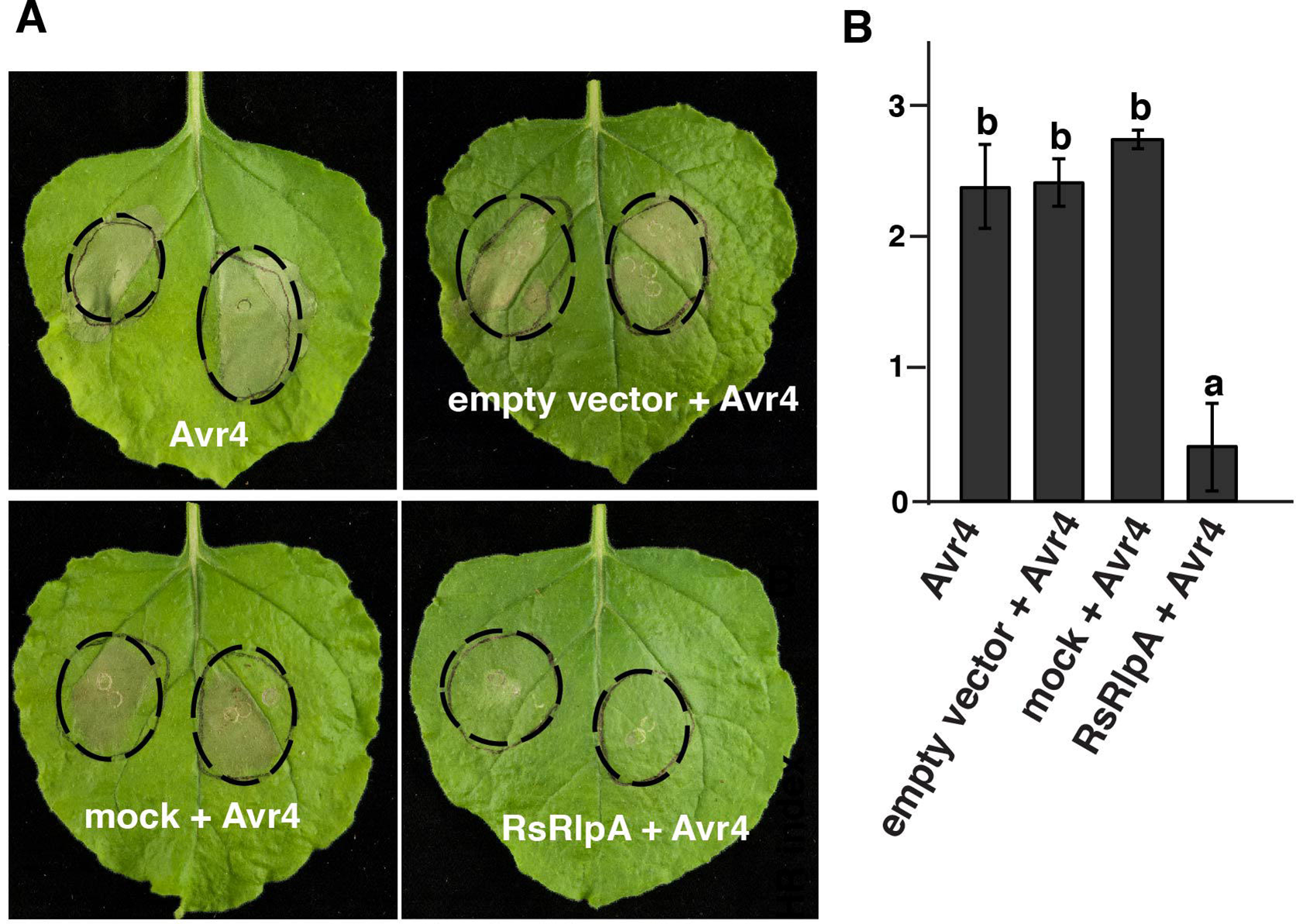
Assay for suppression of Avr4-mediated hypersensitive response (HR). **(A)** Symptoms on transgenic *Nicotiana bethamiana* leaves expressing the tomato Cf-4 receptor. **(B)** HR scale from 0-3 with “0” indicates no symptoms and “3” severe symptoms on *N. bethamiana*. Leaves were agro-infiltrated first with the RsRlpA effector candidate driven by the CaMV:35S promoter and HR challenged 24hpi with the Avr4 effector. Empty vector and mock inoculation were used as controls. Cycles with dashed lines show the agro-infiltration sites of both effectors. Images taken 3dpi. Different letters (a,b) indicate statistical significant differences according to Tukey test (p < 0.05). Error bars represent SD of six plants, each contained two agro-infitrated leaves.

Despite GFP tagging in *RsRlpA+* strains, we repeatedly failed to clearly observe the RsRlpA localization during the infection process in sugar beet leaves. Thus, to determine where this effector is localized, we transiently expressed it in *N. benthamiana* leaves, tagged with GFP or RFP at C-terminus. We observed that RsRlpA was mainly localized to the plasma membrane (**Figure 5**). Localization of RsRlpA was also observed in the nucleoplasm and ER. Plasmolysis of the *N. benthamiana* leaves, using mannitol, supported the plasma membrane localization.

**Figure 5.**
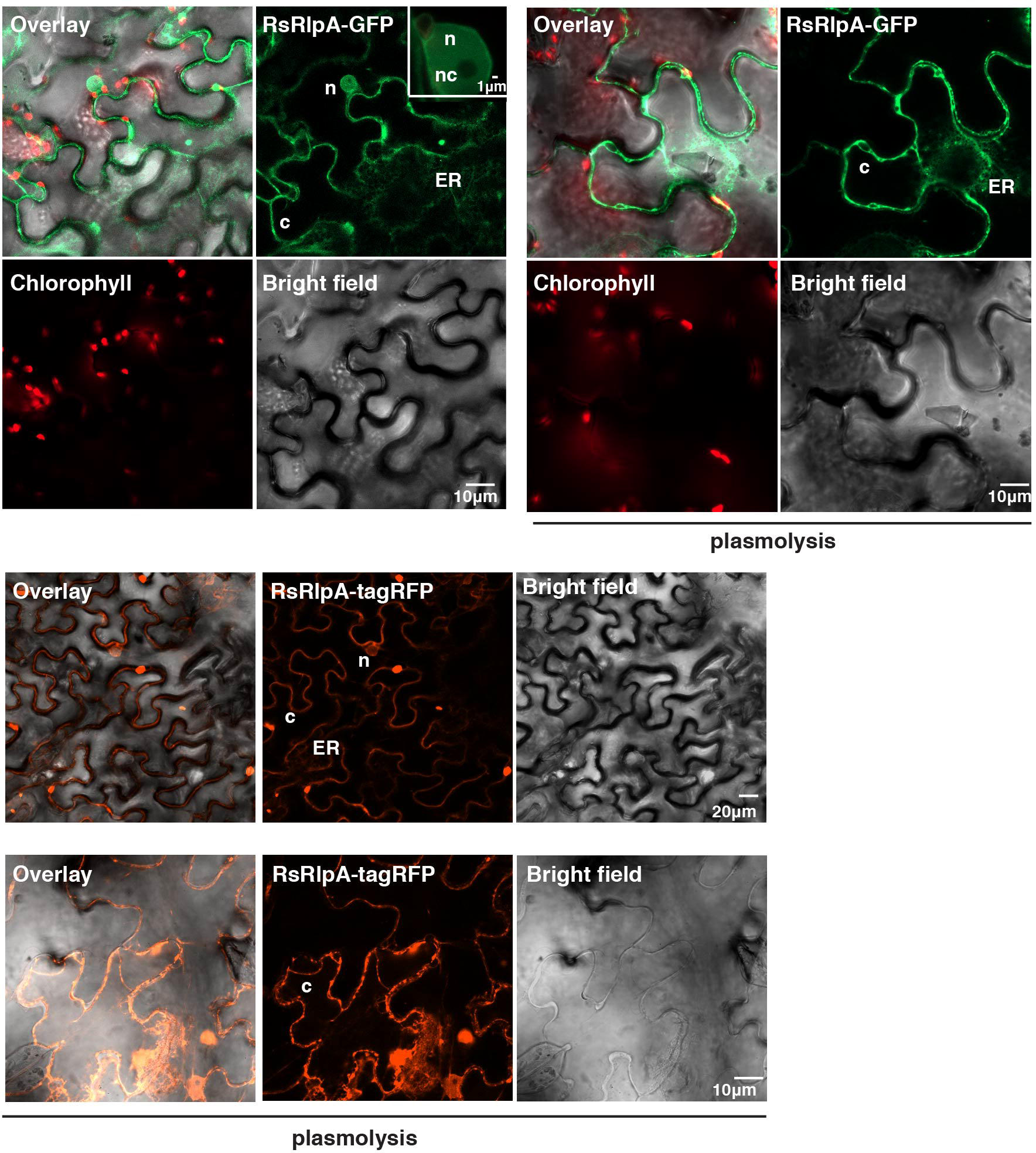
Live-cell imaging of C-terminal GFP-tagged or tagRFP-tagged *R. solani* RsRlpA effector candidate in agro-infiltrated *Nicotiana benthamiana* leaves. The localizations were monitored with a laser-scanning confocal microscope with a sequential scanning mode 48 hrs post infiltration. The GFP and the chlorophyll were excited at 488 nm. GFP (green) and chlorophyll (red) fluorescent signals were collected at 505-525 and 680-700 nm, respectively. The tagRFP was excited at 558 nm and collected at 545-620 nm. Plasmolysis was incited by using 1 M mannitol for 30 min. **(c):** cytoplasm, **(ER):** endoplasmic reticulum, **(n):** nucleoplasm, **(nu):** nucleolus.

### The RsLysM effector binds to chitin

Previous analyses of LysM effector proteins from plant pathogenic ascomycetes have demonstrated their ability to bind chitin (de Jonge *et al.*, 2010; Marshall *et al.*, 2011; Kombrink *et al.*, 2017). Whether LysM effectors from basidiomycetes have a similar chitin-binding affinity is unknown. We expressed *RsLysM* in yeast and purified the protein, which was used for the chitin-binding assay. The *R. solani* LysM protein was able to interact with all tested forms of chitin, such as chitin beads, crab shell chitin and chitosan (**Figure 6A**). No precipitation with other polysaccharides such as xylan and cellulose was detected. Together the data suggests that RsLysM is an active chitin-binding effector protein, similar to other already characterized LysM effectors from filamentous ascomycetes.

**Figure 6.**
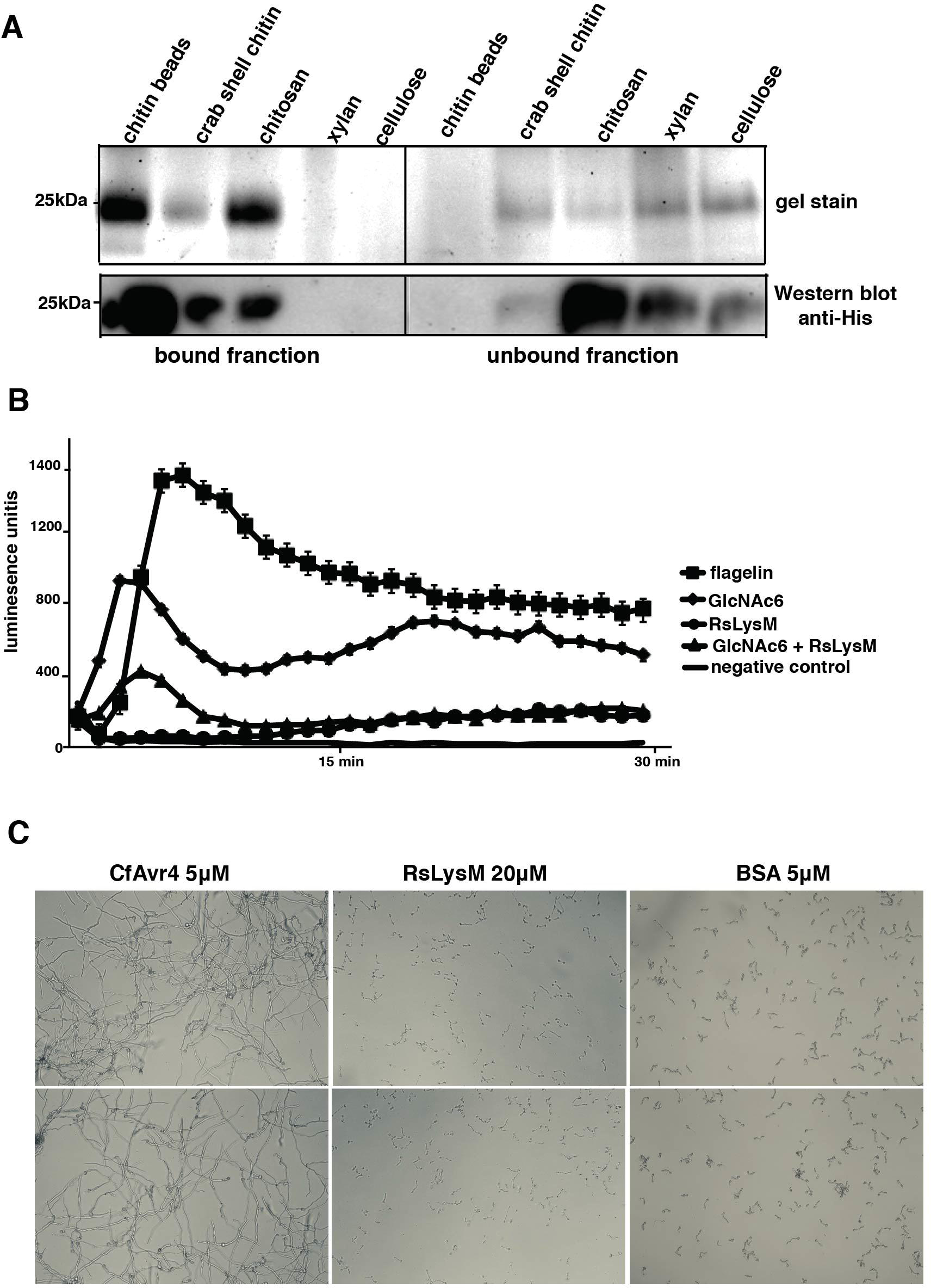
Functional analysis of the RslysM protein. **(A)** Binding affinity assay on different insoluble polysaccharides. 20 μg/ml of LysM protein was mixed with 5 mg of chitin beads, crab shell chitin, chitosan, xylan or cellulose. Presence of protein in bound (pellet) and unbound fraction (supernatant) separated using SDS-polyacrylamide gel electrophoresis followed by Coomassie staining. **(B)** Chitin-induced oxidative (ROS) burst assay in leafs. Production of ROS was determined using luminol-dependent chemiluminescence. Leaf discs were treated with 10nM flagellin (positive control), 10μM GlcNAc6, 10μM LySM protein, 10μM GlcNAc6 + 10μM LySM protein. The luminol reagent was used as a negative control. At least eight biological replicates were run, and experiment repeated twice. **(C)** Hyphal protection assay. 20μM of LysM protein was incubated with 3×10^5^ *T virens* conidia for 30min and 0.1U bacterial chitinase and 1OU zymolyase. Images were taken 6hpi. 5μM Avr4 or BSA were used as positive and negative controls respectively. Assay was repeated twice with same results.

### The RsLysM effector suppresses chitin-triggered immunity but does not protect hyphae from degradation

It is known that LysM effectors stealth filamentous ascomycetes to avoid plant immunity responses triggered by chitin (de Jonge *et al.*, 2010; Marshall *et al.*, 2011; Mentlak *et al.*, 2012; Kombrink *et al.*, 2017). To investigate whether the RsLysM effector also prevent plant chitin-triggered immunity, *N. benthamiana* leaves were treated with chitin oligomers (GlcNAc)_6_, which led to reactive oxygen species (ROS) burst (**Figure 6B**), suggesting that RsLysM displays similar non-recognition function as seen in other pathosystems. Certain LysM effectors are able to protect fungal hyphae from chitinolytic activity (Marshall *et al.*, 2011; Kombrink *et al.*, 2017) similar to the Avr4 effector from *C. fulvum* (van den Burg *et al.*, 2006). To investigate whether the RsLysM displays similar function, germinated conidia from *T. viride* were mixed with pure RsLysM protein, while Avr4 and BSA were used as positive and negative controls respectively. RsLysM was unable to protect fungal hyphae against degradation from bacterial chitinases and zymolyases, in contrast to Avr4 (**Figure 6C**).

## Discussion

*Rhizoctonia solani* is a pathogen commonly described as a saprophyte that is able to switch to pathogenic endotrophic growth thriving on dead or dying plant debris/cells. A complete understanding on the switch of lifestyles from endophytic to a parasitic form is still unclear. Theories underlying the phenomenon are about; an imbalance in nutrient exchange between plant and microorganism, single gene mutations as shown in *Colletotrichum magna*, and light-induced production of H_2_O_2_ (Redman *et al.*, 2001; Alvarez-Loayza *et al.*, 2011; Kuo *et al.*, 2014). Loss of cell wall-degrading enzymes and high activity of certain genes lacking functional annotation has been suggested to at least partly explain the switch to the symbiotic lifestyle of ectomycorrhiza (Kohler *et al.*, 2015). To plan for targeted pathogen control measures mechanisms behind these transitions are essential to be clarified.

The LysM domain is ubiquitous and can be found across all kingdoms. The LysM module recognizes the GlcNAc-X-GlcNAc sequence in polysaccharides (Buist *et al.*, 2008) and details on the binding characteristics are clarified (Mesnage *et al.*, 2014). In prokaryotes, LysM motifs bind to peptidoglycan in bacterial cell walls, whereas LysM in eukaryotes mainly bind to the abundant chitin polymer. In parallel have numerous LysM receptors been identified for example one that binds to nodulation factors important for nodule development in legumes and thereby promote nitrogen fixation (Broghammer *et al.*, 2012). In plant – pathogen interactions, a wealth of knowledge on LysM derives from pathogenic fungal species, (de Jonge *et al.*, 2010; Marshall *et al.*, 2011; Mentlak *et al.*, 2012; Kombrink *et al.*, 2017).

In this study, we investigated the role of two candidate effectors predicted as singletons in the *R. solani* AG2-2IIIB strain (Wibberg *et al.*, 2016b). The RsLysM effector gene showed induction upon early infection stage including suppression of ROS burst, which is in agreement with the previously published data from foliar pathogens such as *C. fulvum* and *Mycosphaerella graminicola* (de Jonge *et al.*, 2010; Marshall *et al.*, 2011). The situation looks different in *Verticillium dahliae*, where a lineage-specific LysM effector contributes to virulence on tomato but not on *N. benthamiana* or *Arabidopsis* (Kombrink *et al.*, 2017).

Our *C. beticola* strains overexpressing the RsLysM displayed increased fungal biomass and necrotic lesions in infected sugar beet plants, strongly supporting its role in host colonization. Similarly, the lineage-specific *Vd2LysM* and Ecp6 effectors are required for full virulence of *V. dahliae* and *C. fulvum* respectively, since deletion of these genes led to reduced symptom development (Bolton *et al.*, 2008; de Jonge *et al.*, 2013). In contrast, deletion of three core LysM effectors from the *V. dahliae* JR2 strain, showed no changes in pathogen virulence (Kombrink *et al.*, 2017), indicating functional differentiation among them. RsLysM did not protect fungal hyphae from degradation, which indicates that it is not involved in protecting the fungal cell wall from external hydrolytic enzymes. This finding is in analogy of the Epc6 LysM effector function (de Jonge *et a*l., 2010). The situation looks different in some other plant-pathogen systems (Marshal *et al.*, 2011; Kombrink *et al.*, 2017), and suggests that hyphal protection is not a universal function for LysM effectors.

The RsRlpA candidate effector is a protein, containing a barwin-like double psi beta-barrel (DPBB) domain. This domain has a possible enzymatic function and it has been identified in various proteins, such as expansins, dehydrogenases, endo-glucanases and proteinases (Castillo *et al.*, 1999). The function of the DPBB domain in filamentous fungi remains unknown, but proteins containing this domain were identified in the secretome analysis of virulent *Pyrenophora tere*s f. *teres* isolates, causing net blotch disease in barley (Ismail and Able, 2016) and in Basidiomycota species causing brown and white rot (Pellegrin *et al.*, 2015). The *RsRlpA* gene was induced at an early stage of sugar beet infection. Induction of a homolog to *RsRlpA* in *Magnaporthe oryzae* is also active in invasive hyphae invaded the first plant cells, indicating a role during the biotrophic stage of this hemibiotrophic pathogen (Mosquera *et al.*, 2009). The *C. beticola* RsRlpA overexpression strains, displayed increased fungal biomass in infected plants, and to our best of knowledge, this is the first report showing the contribution of this kind of protein in fungal virulence.

Although more studies are needed in order to characterize the RsRlpA enzymatic function and the HR suppression, this study gave us valuable information about the potential role of effectors in the AG2-2IIIB *R. solani* strain. It seems that this pathogen deploys effectors to manipulate important plant immunity mechanisms and it does not rely only to necrosis-induced effectors. We could also speculate that *R. solani* AG2-2IIIB needs an initial biotrophic stage crucial for a successful establishment of infection, implying a hemibiotrophic life style.

## Supplementary material

**Figure S1**. Validation of *C. beticola* overexpression strains harboring the *R. solani RsLySM* and *RsRlpA* genes. Transcript from cDNA extracted from mycelia grown on PDA medium, for 7days. *C. beticola* wild type (WT) was used as negative control and *act* gene used as a positive control.

**Tables S1**. Primers used in the current study

## Acknowledgments

The authors want to acknowledge: Prof. Bart Thomma, Peter Herfs, Nick Snelders and Hui Tian at Wageningen University, Phytopathology Lab, who contributed to RsLysM protein production, purification and provided us with *N. benthamiana* Cf-4 seeds, MariboHilleshög Research that provided *C. beticola* Ty1 strain and sugar beet seeds (breeding line 16045118 01), to Dr. Ioannis Stergiopoulos and Dr. Li Hung-Chen at UC Davis, Phytopathology Lab for helping us with the hyphal protection assay. This work was supported by grants from: the Swedish Research Councils VR and Formas, MariboHilleshög Research and the Swedish University of Agricultural Sciences.

